# Asparagine availability differentially regulates early vs late CD4^+^ and CD8^+^ T cell activation, metabolism and autophagy

**DOI:** 10.64898/2026.04.27.721062

**Authors:** Mingcheng Song, Linda V. Sinclair, Mary Tozer, Mihaela Lorger, Robert J. Salmond

## Abstract

T cell activation is associated with, and dependent upon, the upregulation of amino acid uptake from the extracellular environment. Uptake of the non-essential amino acid asparagine (Asn) is mediated via amino transporters such as Slc1a5 whilst Asn can be synthesized within cells that express asparagine synthetase (ASNS). Previous work demonstrated that initial activation of CD8^+^ T cells is perturbed in the absence of Asn, whereas effector cytotoxic T cells cells upregulate ASNS and lose their dependence on Asn uptake. By contrast, less is known of the role of Asn uptake and ASNS in CD4^+^ T cell responses. Here we demonstrate that CD4^+^ T cells are more reliant than CD8^+^ T cells on Asn uptake for initial activation, differentiation, metabolic reprogramming and regulation of autophagy. These phenotypes are associated with enhanced expression of ASNS in CD8^+^ as compared to CD4^+^ effector T cells.

## Introduction

T cells integrate antigen, costimulatory and cytokine receptor signalling to facilitate activation and effector responses. These activatory signals converge on a core set of signalling pathways, including but not limited to mechanistic target of rapamycin (mTOR) [1; 2] and Myc-dependent pathways [3; 4], to enable metabolic reprogramming. This results in a switch from a basal catabolic metabolism, typical of naïve T cells, to anabolism and thereby provides the energy and cellular building blocks to sustain T cell growth, proliferation, differentiation and effector responses [5]. A major consequence of and prerequisite for T cell activation is the upregulation of cell surface nutrient transporter expression, that enables enhanced nutrient uptake to fuel ongoing immune responses. Thus, genetic deletion of individual glucose [6; 7; 8] or amino acid transporters [9; 10] results in defective T cell responses. Similarly, low nutrient availability *in vivo*, for example in the tumour microenvironment, is linked to defective T cell activation and responses [11; 12]. Glucose uptake from the environment is required to fuel T cell glycolysis which provides ATP as well as precursors for biosynthetic pathways [2]. Amino acid uptake is required for T cell protein synthesis, fuels the TCA cycle, regulates mTOR signalling pathways, and contributes to numerous biosynthetic pathways and the regulation of cellular redox state. Recent studies have also shown a key role for amino acid uptake and availability for the repression of autophagy in activated and effector CD8^+^ T cells [13].

Despite a capacity for their intracellular synthesis, the uptake of several non-essential amino acids such as glutamine, arginine, serine and glycine is required for optimal T cell activation [14]. Asparagine (Asn) is a non-essential amino acid that can be synthesised within cells that express asparagine synthetase (ASNS). Previous studies determined that naïve CD8^+^ T cells express low levels of ASNS [15] and consequently early stages of CD8^+^ T cell activation require the uptake of Asn [15; 16; 17] from the environment, likely via the Slc1a5/ASCT2 amino acid transporter [18; 19]. During activation, CD8^+^ T cells upregulate ASNS that enables effector cells to function in Asn-depleted conditions [15; 17; 20]. Asn is mainly required for protein synthesis but may also contribute to the regulation of T cell signalling via effects on Lck and mTOR pathways [15; 16]. Activation in Asn-deprived conditions initially impedes T cell responses, however can result, in the longer term, in enhanced T cell metabolic fitness and functionality as a consequence of metabolic rewiring [20]. Indeed, a recent study reported that treatment of a small cohort of patients with asparaginases, that degrade Asn, enhances T cell activity in metastatic nasopharyngeal carcinoma and improves responses to immune checkpoint blockade [21]. In contrast, several studies have reported an inhibitory effect of asparaginases on CD8^+^ T cell activation [22; 23].

Studies on the role of Asn metabolism in T cells have thus far focused on CD8^+^ T cells, as the main effector population in cancer. In the current work, we compared directly the CD4^+^ and CD8^+^ T cell requirement for extracellular Asn. We report that, in comparison to CD8^+^ T cells, CD4^+^ T cells are more reliant on Asn uptake for initial activation, differentiation, metabolic reprogramming and regulation of autophagy, whilst this is linked to enhanced levels of ASNS expression in effector CD8^+^, as compared to CD4^+^, T cell subsets.

## Results

### Asn uptake and ASNS expression are required for optimal CD4^+^ T cell activation

Our previous studies determined that, upon activation, CD8^+^ T cells coordinate Asn uptake with intracellular Asn biosynthesis via ASNS [15]. To determine whether CD4^+^ T cells had similar requirements, purified C57BL/6 naïve CD4^+^ T cells were activated for 48h under TH1-polarizing conditions (anti-CD3/CD28 + IL-12) in DMEM culture media supplemented ± 300 μM Asn ± 2 mM Gln. Flow cytometry analysis of live/dead staining showed that proportions of live CD4^+^ T cells were substantially reduced following activation in Asn-and, particularly, Gln-free conditions (Figure 1A). TCR-induced upregulation of the Myc target CD71 (transferrin receptor) (Figure 1B-C) and the Th1-associated transcription factor Tbet (Figure 1E-F) was completely blocked in the absence of extracellular Gln. Proportions of CD71^+^ and Tbet^+^ CD4^+^ T cells were reduced in the absence of Asn but not completely blocked. Analysis of geometric mean fluorescence intensities of marker staining in gated positive cells demonstrated that levels of CD71 (Figure 1D) and Tbet (Figure 1G) expression were reduced in Asn-free conditions. Furthermore, TCR-induced CD4^+^ T cell IL-2 production was also severely impeded under conditions of Asn or Gln deprivation (Figure 1H).

**Figure 1.**
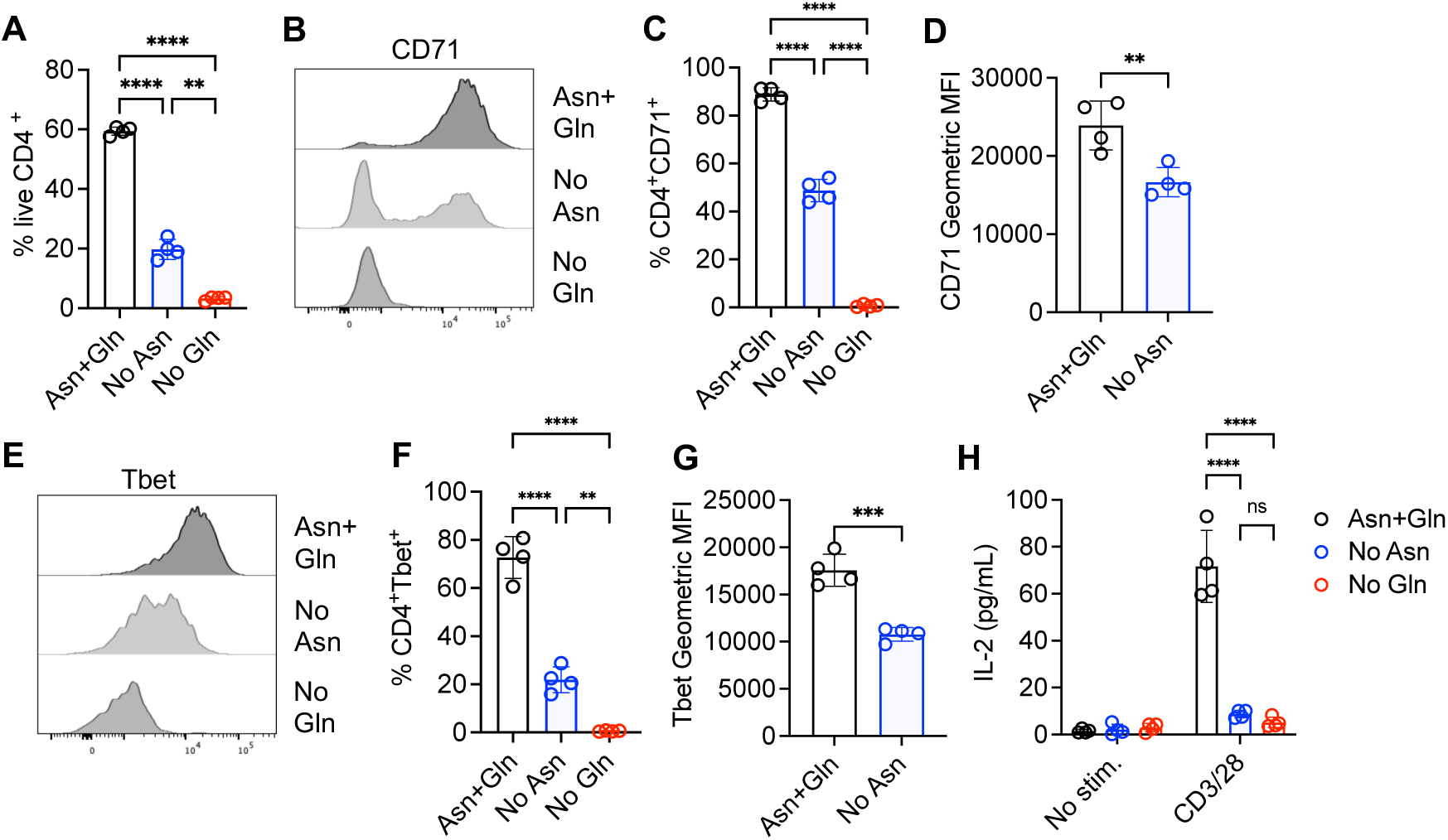
Asn availability is required for optimal CD4^+^ T cell activation. Purified naïve C57BL/6 CD4^+^ T cells were stimulated with CD3 / CD28 antibodies (A-H) and IL-12 (A-G) in DMEM culture media supplemented ± 300μM Asn and 2mM Gln for 48h (A-G) or 24h (H). Proportions of live T cells were determined by live-dead staining and flow cytometry (A). Surface CD71 (B) and intracellular Tbet (E) expression on live CD4^+^ T cells were assessed by flow cytometry. Graphs represent % CD71^+^ (C) or Tbet^+^ (F) CD4^+^ T cells or geometric mean fluorescence intensity (MFI) values from gated CD71^+^ (D) or Tbet^+^ (G) cells. Levels of IL-2 in culture supernatants were assessed by ELISA (H). In all cases, dots represent biological replicates (n=4) from 1 of 3 repeated experiments. Bars represent mean values of replicate samples and error bars represent SD. ns – not significant, * p<0.05, ** p<0.01, *** p<0.001, **** p<0.0001 as determined by one-way (A, C, F) or two-way ANOVA (H) with Tukey’s multiple comparisons test or unpaired Student’s t-test (D, G).

### CD4^+^ T cell requirement for Asn is more stringent than CD8^+^ T cells’

Together, our analysis of purified CD4^+^ T cells (Figure 1) and our previous studies of OT-I TCR transgenic T cells [15] suggested that CD4^+^ and CD8^+^ T cells both require extracellular sources of Asn during initial activation. To compare the two cell types directly, we performed activation experiments using unsorted polyclonal lymph node T cells. LN cells were activated under TH1-polarizing conditions in DMEM medium ± 300μM Asn and expression of CD71 and Tbet assessed by flow cytometry at 48h.

Following activation in Asn-replete media, ∼90% of both CD4^+^ and CD8^+^ T cells had upregulated CD71 expression, with a majority of activated CD71^+^ cells also expressing Tbet (Figures 2A-C). Stimulation under conditions of Asn-deprivation limited the proportions of activated CD71^+^Tbet^+^ CD4^+^ and CD8^+^ T cells, however the requirement for Asn was more stringent for CD4^+^ populations. Thus, ∼15-20% of CD4^+^ T cells had upregulated both CD71 and Tbet following 48h of activation in Asn-free media as compared to ∼35-45% of CD8^+^ T cells (Figure 2A). Titration experiments demonstrated that at Asn concentrations ≥33 μM, CD4^+^ and CD8^+^ T cells were equally and optimally activated in TH1-polarizing conditions, as assessed by proportions of CD71^+^ (Figure 2D) and Tbet^+^ cells (Figure 2E). By contrast, while the activation of both CD4^+^ and CD8^+^ T cells was impeded at lower Asn concentrations, CD4^+^ T cells were more affected than CD8^+^ T cells (Figure 2D, 2E). Therefore, Asn uptake is required for optimal activation of both CD4^+^ and CD8^+^ T cells, but CD8^+^ T cells have greater capacity for activation when Asn is limiting.

**Figure 2.**
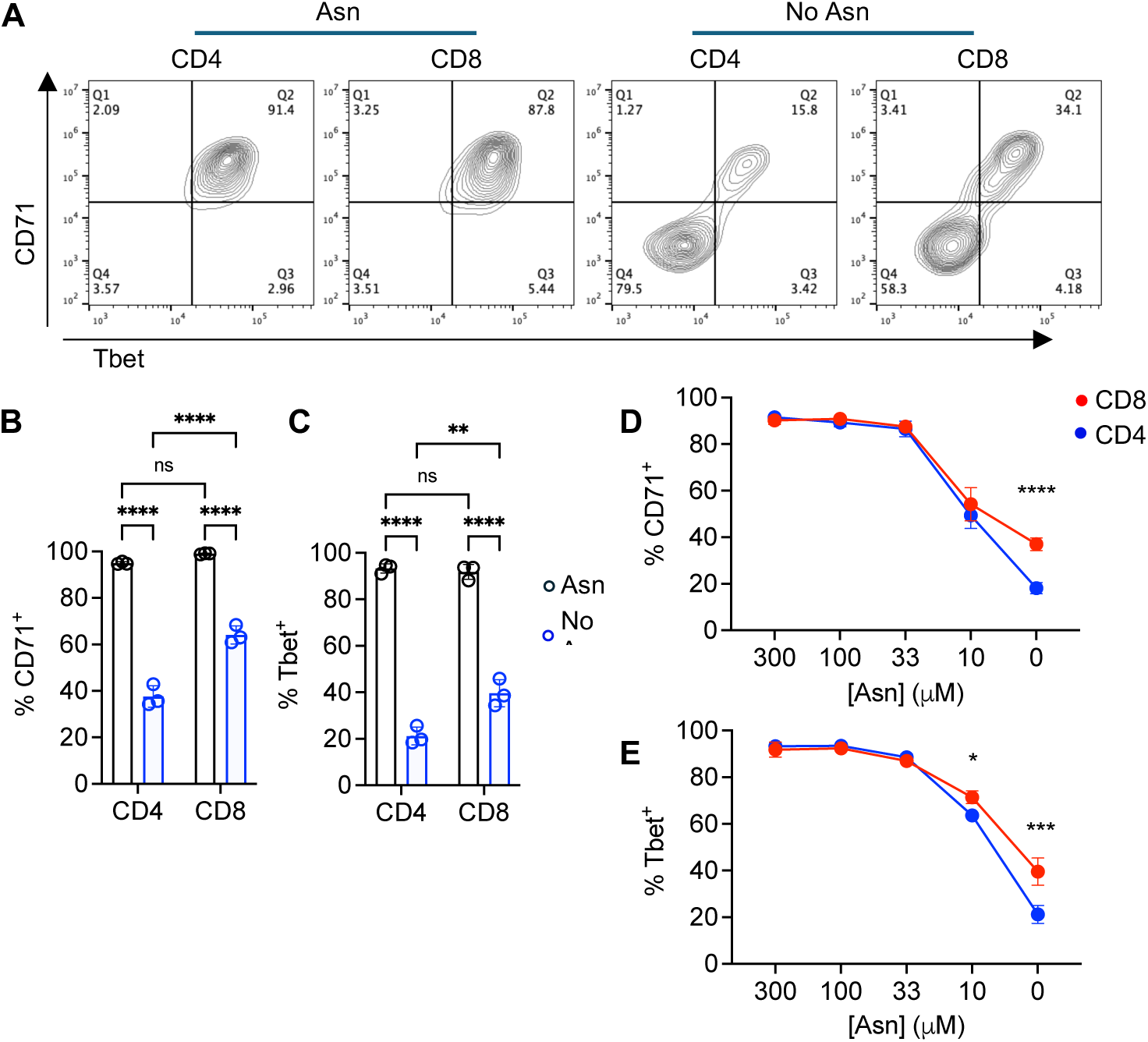
CD4^+^ T cell requirement for Asn is more stringent than CD8^+^ T cells’. Lymph node T cells were stimulated for 48h with CD3 / CD28 antibodies and IL-12 in DMEM culture media supplemented ± Asn. (A) Representative dotplots of gated live CD4^+^ and CD8^+^ T cell CD71 and Tbet expression. Values in dotplots represent proportions of cells within indicated quadrants. Surface CD71 (B) and intracellular Tbet (C) expression was assessed by flow cytometry. Dots represent biological replicates (N-3) from 1 of 4 repeated experiments. Titration of Asn levels demonstrated enhanced capacity for CD8^+^ T cell upregulation of CD71 (D) and Tbet (E) in Asn-limiting conditions. Data are from 1 of 3 repeated experiments. In all cases, error bars represent SD. ns – not significant, * p<0.05, ** p<0.01, *** p<0.001, **** p<0.0001 as determined by two-way ANOVA with Tukey’s multiple comparisons test.

### Asparagine is required for optimal differentiation of effector cytokine-producing T cells

Recent studies have reported that asparagine-deprivation limits initial activation but may confer CD8^+^ T cells with enhanced effector capacity [20; 21]. To further analyse these phenotypes for both CD4^+^ and CD8^+^ T cells, LN cells were activated for 72 h in TH1-polarizing conditions ± 300 μM Asn. Cells were then transferred to Asn-replete conditions and re-stimulated with PDBU/ionomycin to maximally induce effector cytokine production. Results demonstrated that when LN cells had been initially activated in the presence of Asn, ∼40% of CD4^+^ and ∼90% of CD8^+^ were competent to produce IFNγ, whilst ∼60% of both CD4^+^ and CD8^+^ T cells produced TNF, as assessed by flow cytometry (Figure 3). Strikingly, <5% of CD4^+^ T cells initially activated in the absence of Asn were capable of producing IFNγ. CD8^+^ T cell capacity for IFNγ production was also affected, however ∼30% of the cells initially activated in the absence of Asn produced IFNγ upon restimulation (Figure 3A, 3B). Further analysis demonstrated that the proportions of CD4^+^ T cells and CD8^+^ T cells competent to produce IFNγ after initial activation in the presence as compared to the absence of Asn was increased ∼12-fold and ∼3-fold, respectively (Figure 3C), demonstrating the increased reliance of CD4^+^ T cells for Asn uptake. Despite this, levels of IFNγ production per cell, as assessed by analysis of geometric mean fluorescence intensities of gated cytokine cells, was reduced for CD8^+^ but not CD4^+^ T cells activated in the absence of Asn (Figure 3D). By contrast, the capacity for both CD4^+^ and CD8^+^ T cells to produce TNF was only mildly impacted by initial activation under conditions of Asn deprivation (Figure 3A, 3E, 3F).

**Figure 3.**
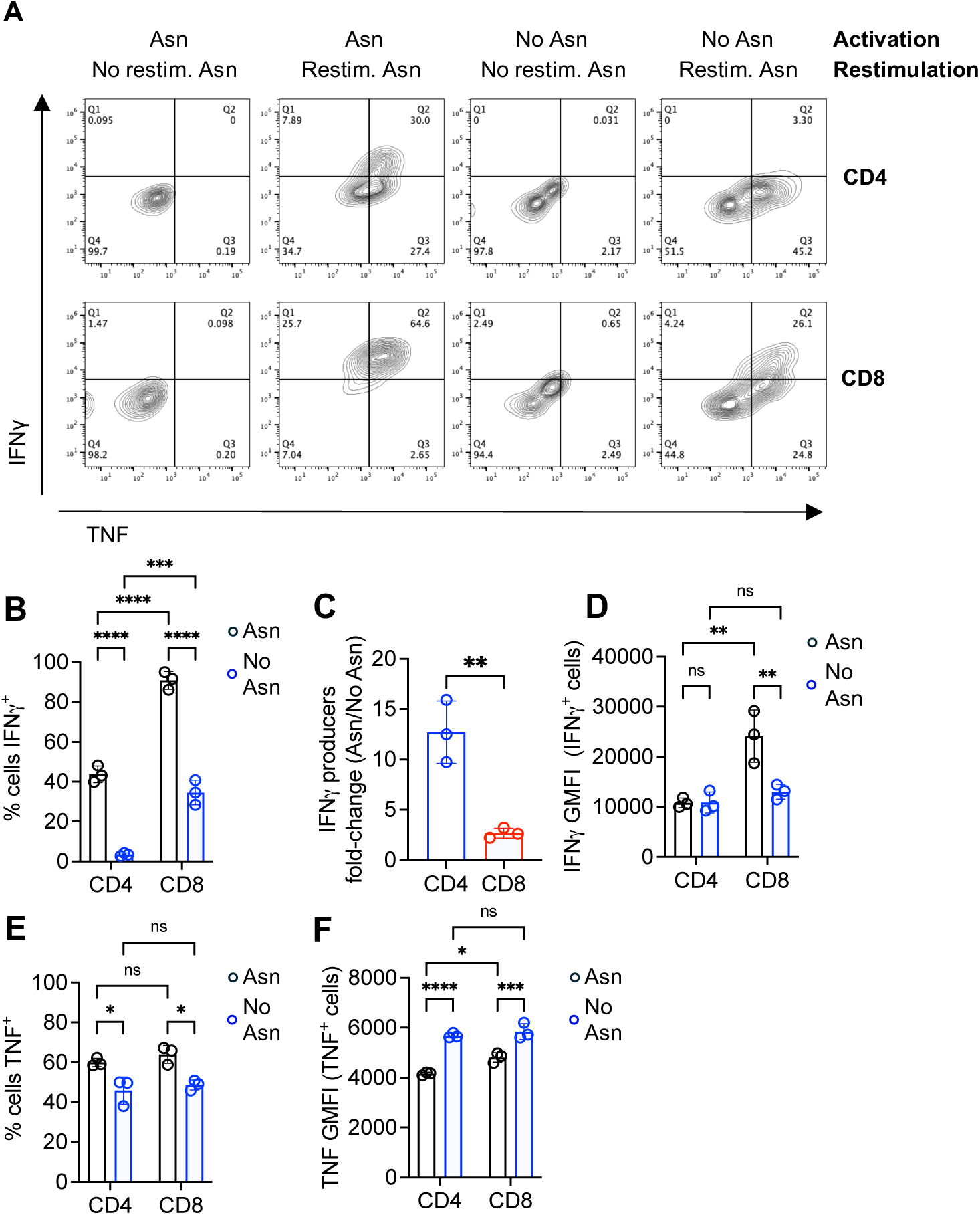
Differentiation of IFNγ-producing T cells is impeded in the absence of Asn. Lymph node T cells were stimulated for 72h with CD3 / CD28 antibodies and IL-12 in DMEM culture media supplemented ± 300μM Asn, then restimulated with PDBU/Ionomycin in Asn-replete media. Representative dotplots (A) show levels of IFNγ and TNF staining in gated CD4^+^ and CD8^+^ T cells, whilst barcharts show mean proportions of IFNγ^+^ (B) and TNF^+^ (E) cells or geometric mean fluorescence intensity (GMFI) of cytokine staining in gated cytokine-positive cells (D, F). (C) Barchart shows fold-change increase in proportions of IFNγ-producing CD4^+^ and CD8^+^ T cells activated in Asn-containing conditions as compared to Asn-free media. Data are from 1 of 3 repeated experiments. In all cases, error bars represent SD. ns – not significant, * p<0.05, ** p<0.01, *** p<0.001, **** p<0.0001 as determined by two-way ANOVA with Tukey’s multiple comparisons test.

Effector T cells may gain the capacity to function independently of extracellular Asn [15]. To compare directly the impact of Asn availability on effector CD4^+^ and CD8^+^ T cell cytokine production, LN cells were activated for 72 h in TH1-polarizing conditions in the presence of Asn, followed by re-stimulation with PDBU/ionomycin ± Asn. Depletion of Asn solely during the re-stimulation stage did not impact the capacity of effector CD4^+^ or CD8^+^ T cells to produce either IFNγ or TNF (Supplementary Figure 1). Therefore, Asn availability is limiting for the differentiation of both CD4^+^ and CD8^+^ T cells to an IFNγ-producing phenotype, with CD4^+^ T cells more markedly impacted than CD8^+^ T cells. However, once T cells have undergone Type 1 differentiation, Asn is no longer required for effector responses.

### CD8^+^ T cells have greater capacity than CD4^+^ T cells for activation-induced metabolic reprogramming in the absence of Asn

To link effects of Asn availability on T cell activation to metabolic phenotypes, we used BioTracker-ATP^TM^ (BT-ATP), a fluorescent probe that specifically reports mitochondrial ATP levels in live cells. LN T cells were activated for 48h in TH1 conditions ± Asn, then were co-stained with CD4, CD8 and CD71 mAbs and BT-ATP. When activated in the presence of Asn, both CD4^+^ and CD8^+^ T cells uniformly expressed CD71 and demonstrated high levels of BT-ATP staining (Figures 4A, 4B). In the absence of Asn, proportions of CD71^+^BT-ATP^hi^ cells were reduced, with CD4^+^ T cells (∼28%) being more affected than CD8^+^ T cells (∼50%, Figure 4B). However, when analysis was performed on gated CD71^+^ cells, BT-ATP mean fluorescence intensities were comparable for CD4^+^ and CD8^+^ T cells activated in the absence or presence of Asn (Figure 4C). Therefore, Asn availability limits the proportions of T cells capable of becoming activated and upregulating anabolic metabolism, with CD4^+^ T cells more affected than CD8^+^ T cells. However, in T cells which overcome the activation threshold in Asn-deprived conditions, the levels of metabolic activity, as judged by ATP production, are similar to that of control cells in Asn-replete conditions.

**Figure 4.**
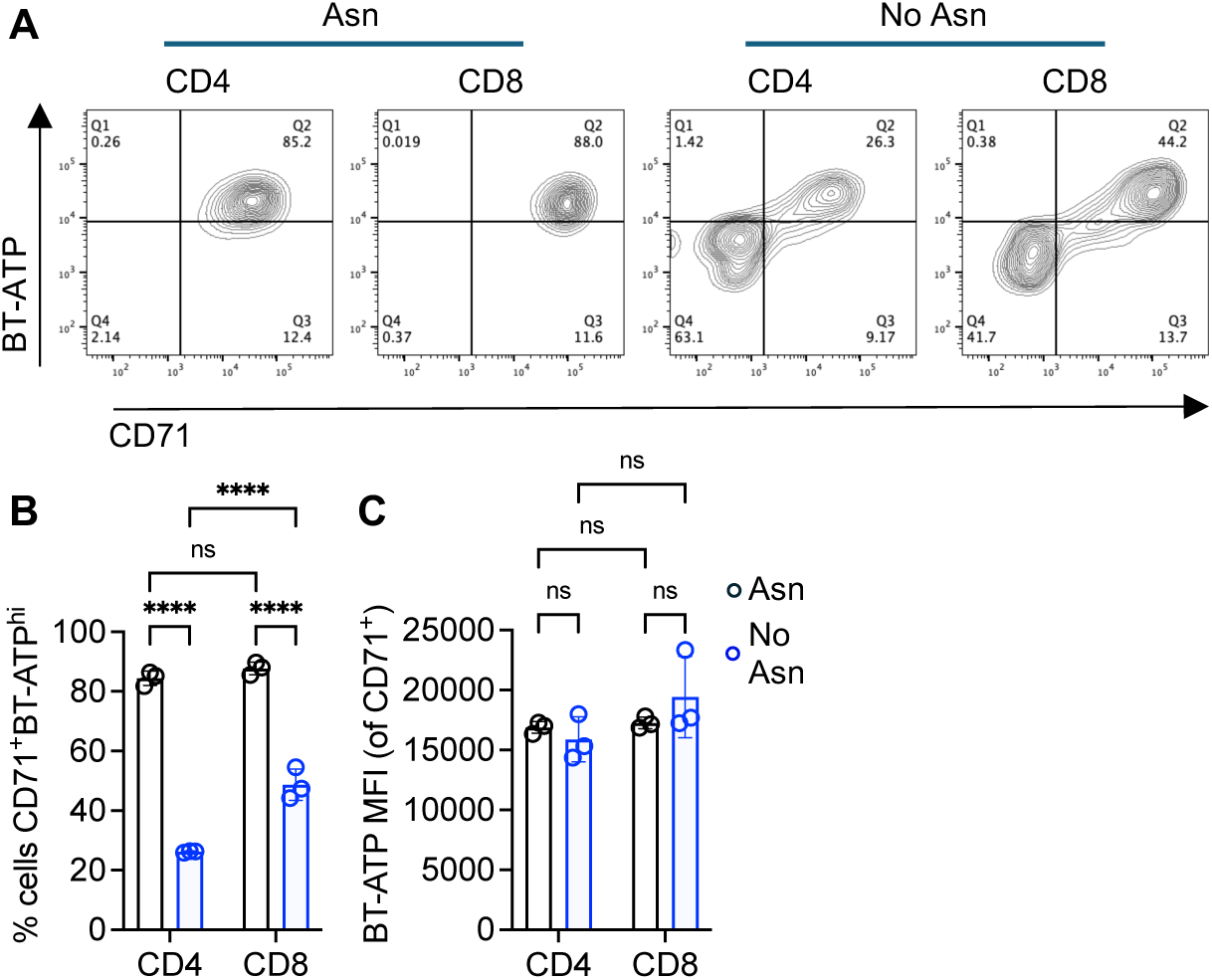
Asn availability is limiting for activation-induced upregulation of anabolic metabolism. Lymph node T cells were stimulated for 48h (A-C) with CD3 / CD28 antibodies and IL-12 in DMEM culture media supplemented ± Asn. Representative dotplots (A) show levels of CD71 and BioTracker-ATP staining in gated CD4^+^ and CD8^+^ T cells, whilst barcharts show mean proportions of CD71^+^BT-ATP^high^ cells (B) and BT-ATP mean fluorescence intensity on gated CD71^+^ cells (C). Data are from 1 of 3 repeated experiments. In all cases, dots represent replicate values and error bars represent SD. ns – not significant, **** p<0.0001 as determined by two-way ANOVA with Tukey’s multiple comparisons test.

### Asparagine deprivation differentially influences autophagy during early T cell activation versus effector T cells

Autophagy is repressed in T cells by antigen and cytokine signals as a consequence of upregulated amino acid transporter expression and amino acid uptake [13]. Conversely, autophagy is reintroduced in effector T cells under conditions of amino acid starvation. We sought to determine the role of Asn availability in the dynamic regulation of T cell autophagy using an autophagic flux reporter. In this mouse strain, a dual fluorescent-tagged LC3b protein (mCherry-GFP-LC3b) is expressed ubiquitously, and upon autophagy induction is recruited into autolysosomes, where low pH quenches GFP fluorescence, with no effect on mCherry. Therefore, autophagic flux can be assessed by flow cytometry analysis of levels of GFP vs mCherry fluorescence; no autophagy – GFP vs mCherry levels are linear, autophagic flux – quenching of GFP signal vs mCherry (Supplementary Figure 2A). LN T cells were activated under TH1-polarizing conditions ± 300 μM Asn for 18h and autophagic flux assessed. Control CD4^+^ and CD8^+^ T cells maintained in the presence of the homeostatic cytokine IL-7 had high levels of autophagy, as demonstrated by low GFP vs mCherry LC3b signal whereas TCR-stimulated T cells substantially downregulated autophagic flux (Supplementary Figure 2B, Figures 5A, B). Importantly, the repression of autophagy following T cell activation was much impeded under conditions of Asn deprivation with a more striking effect on CD4^+^ (Figure 5A) as compared to CD8^+^ T cells (Figure 5B).

**Figure 5.**
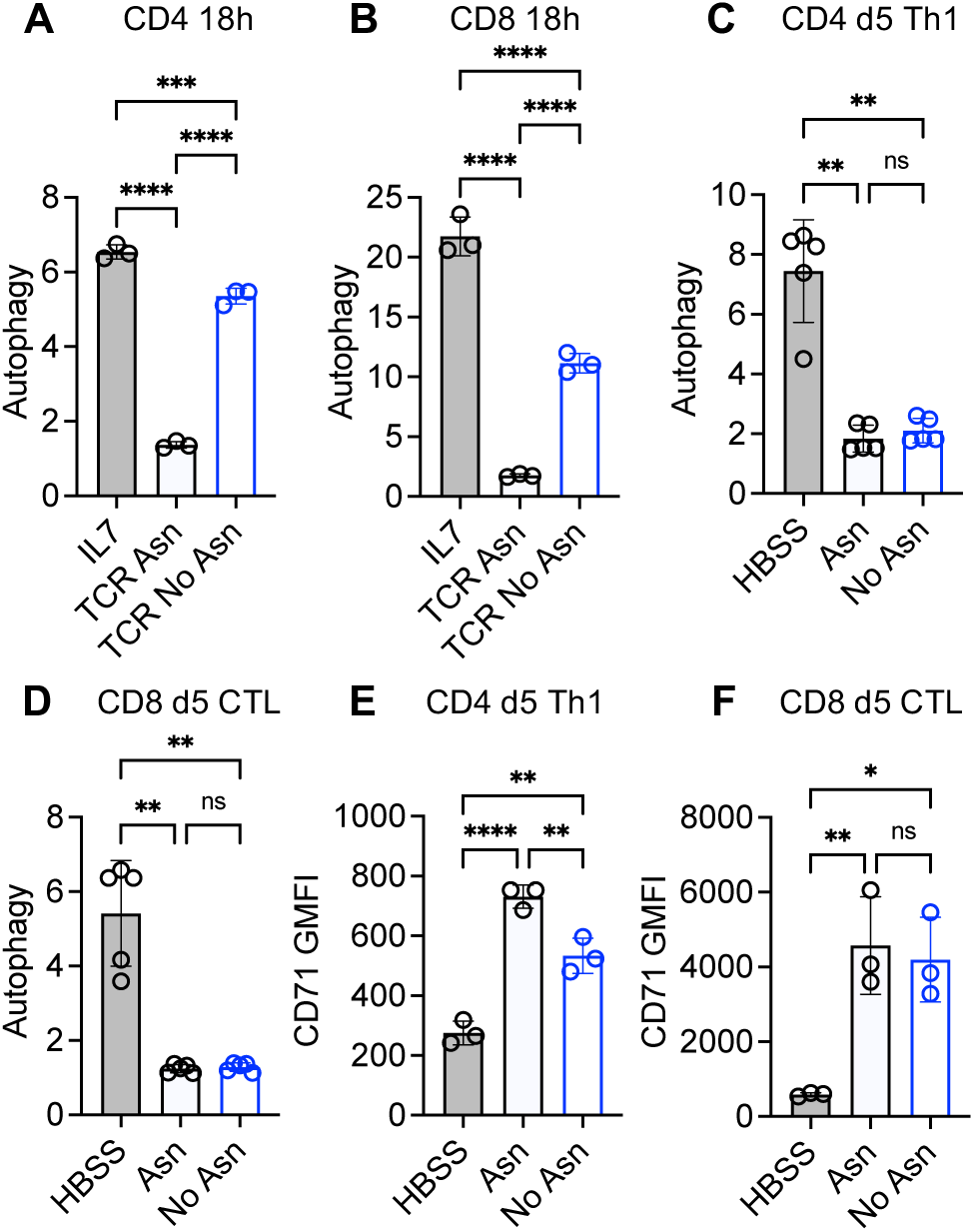
Asn availability regulates activation-induced repression of autophagy in naïve T cells but is not required for suppression of autophagy in effector T cells. Levels of autophagic flux were assessed using mCherry-GFP-LC3b reporter mice. (A, B) Lymph node T cells from autophagy reporter mice were stimulated for 18h with CD3 / CD28 antibodies and IL-12 in DMEM culture media supplemented ± 300 μM Asn. Barcharts show levels of autophagy in gated CD4^+^ (A) and CD8^+^ (B) cells. (C-F) Effector T cells from autophagy reporter mice were generated by 5 days of stimulation *in vitro*. Cells were switched into DMEM ± 300 μM Asn or amino acid-free conditions (HBSS) for 2h followed by assessment of autophagy and CD71 expression by flow cytometry. Bar charts show levels of autophagy and CD71 geometric mean fluorescence intensity (GMFI) in gated CD4^+^ (C, E) and CD8^+^ (D, F) cells. In all cases, dots represent biological replicate values (n=3-5) and error bars represent SD. ns – not significant, * p<0.05, *** p<0.001, **** p<0.0001 as determined by one-way ANOVA (B, C, G, H) with Tukey’s multiple comparisons test or Brown-Forsythe and Welch ANOVA (E, F) with Dunnett’s T3 multiple comparisons test.

To test the role of Asn availability in suppression of autophagic flux in effector cells, LN T cells were activated under TH1-polarising conditions in for 5 days, then were switched to DMEM media ± Asn, or HBSS. In effector CD4^+^ and CD8^+^ T cells maintained in DMEM with 300 μM Asn, autophagy levels were low (Supplementary Fig. 2C, Figures 5C, D). Switching effector cells to amino acid-free conditions (HBSS) reintroduced autophagic flux as demonstrated by GFP quenching, whereas switching to Asn-free DMEM had no effect on autophagy for both CD4^+^ (Figure 5C) and CD8^+^ (Figure 5D) cells. Therefore, Asn availability affects the capacity of naïve T cells to downregulate autophagy following activation but does not affect autophagy in effector cells. Despite not affecting autophagic flux, switching to Asn-free conditions resulted in a reduction in CD71 expression by effector CD4^+^ (Figure 5E) but not CD8^+^ T cells (Figure 5F).

### Enhanced capacity for CD8^+^ T cell activation in absence of Asn is associated with elevated ASNS expression

We previously determined that ASNS expression was required for CD8^+^ T cell activation under conditions of Asn-deprivation [15]. To assess directly the role of ASNS expression in CD4^+^ T cell activation, the *Asns*^Tm1a^ gene-trap mouse was used. In this model, homozygous expression of a hypomorphic *Asns* allele results in a >90% reduction in ASNS protein expression in T cells [15]. Notably, CD4^+^ T cell activation, as assessed by analysis of TCR-induced CD71 expression, was completely blocked in the combined absence of ASNS and extracellular Asn (Figure 6A), similar to our previously reported results assessing CD8^+^ T cell activation [15].

**Figure 6.**
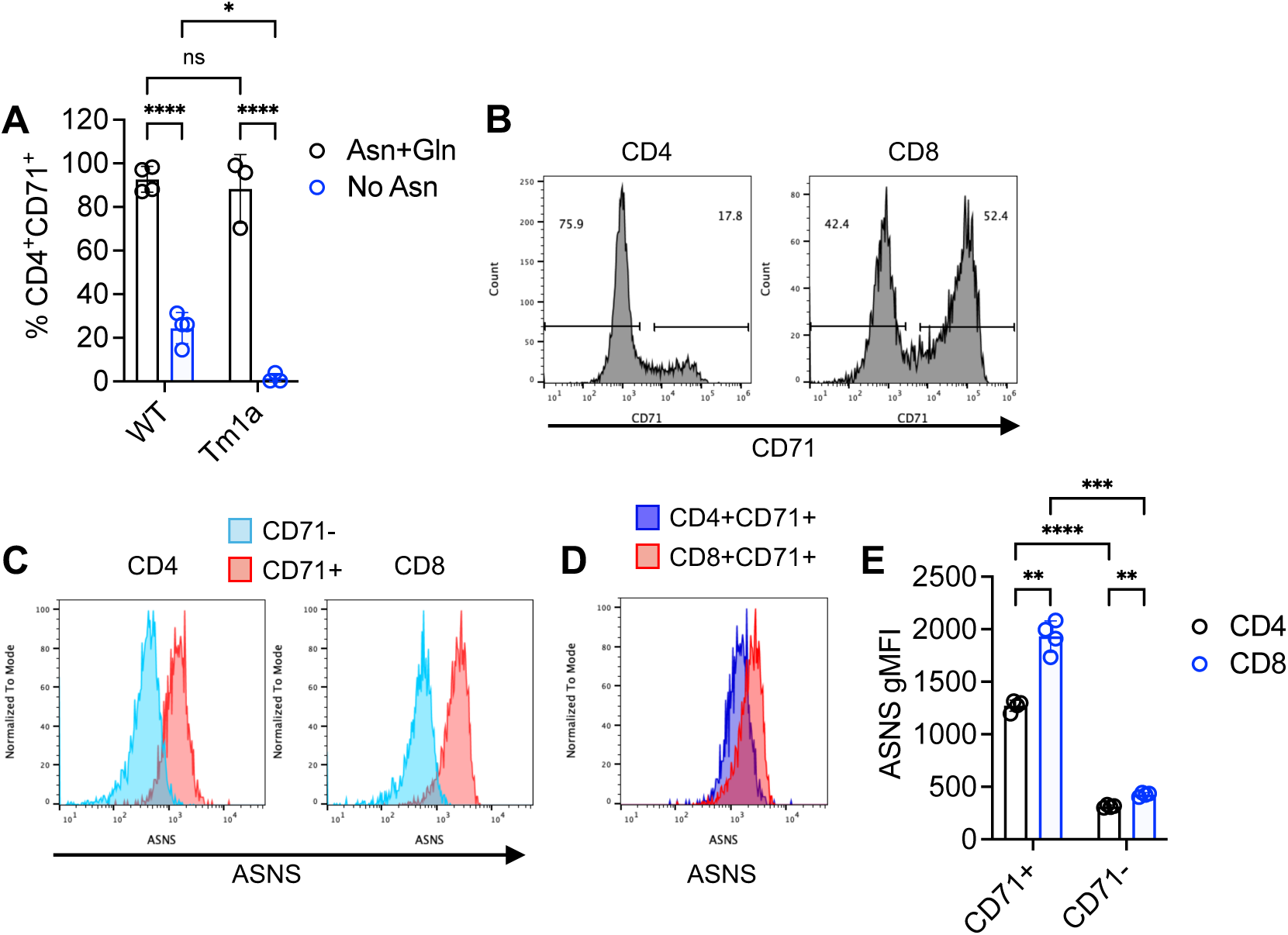
Enhanced capacity for CD8^+^ T cell activation in absence of Asn is associated with elevated ASNS expression. Lymph node T cells from *Asns*^WT/WT^ or *Asns*^Tm1a/Tm1a^ mice (A) or C57BL/6 mice (B-E) or were stimulated for 48h with CD3 / CD28 antibodies in DMEM culture media ± Asn (A) or with no Asn (B-E). (A) Barchart shows proportions of CD4^+^CD71^+^ cells. (B) Representative histograms show levels of CD71 in gated CD4^+^ and CD8^+^ T cells. (C) Expression of CD71 under conditions of Asn deprivation was associated with elevated levels of ASNS expression. (D) Histogram shows that activated CD8+ T cells express higher levels of ASNS than activated CD4+ T cells. (E) Barchart show geometric MFI of ASNS expression in gated CD71^+^ and CD71^-^ T cells. In all cases, circles represent biological replicates (n=3-4). ns – not significant, * p<0.05, ** p<0.01, *** p<0.001, **** p<0.0001 as determined by two-way ANOVA with Tukey’s multiple comparisons test.

As ASNS was functionally implicated in the activation of both CD4^+^ and CD8^+^ T cells, we sought to determined whether ASNS expression levels underpin the lower reliance of CD8^+^ as compared to CD4^+^ T cells on Asn uptake. To compare the regulation of ASNS expression in CD4^+^ and CD8^+^ T cells under conditions of Asn deprivation, we activated LN cells for 48h with CD3/CD28 antibodies. As before, proportions of CD71^+^ T cells were reduced for CD4^+^ T cells as compared to CD8^+^ T cells (Figure 6B). Intracellular staining demonstrated that ASNS expression was ∼3-4 fold higher in activated CD71^+^ as compared to CD71^-^ cells in both CD4^+^ and CD8^+^ populations (Figure 6C), consistent with ASNS being associated with or required for T cell activation in these conditions. Of note, activated CD8^+^CD71^+^ T cells expressed significantly higher levels of ASNS as compared to CD4^+^CD71^+^ T cells (Figure 6D, 6E). Furthermore, proteomic data from the Immunological Proteome Resource [24] demonstrated that ASNS protein copy number is uniformly low in naïve T cells but substantially higher in effector CD8^+^ T cell populations as compared to equivalent CD4^+^ T cells (Supplementary Figure 3). Therefore, CD8^+^ T cells may be more resilient than CD4^+^ T cells under conditions of Asn-deprivation due to an enhanced capacity for intracellular Asn synthesis.

## Discussion

The availability and uptake of several non-essential amino acids is required for the activation, differentiation and effector capacity of T cells. In the current work, we compared the requirements of CD4^+^ and CD8^+^ T cells for extracellular Asn during activation, differentiation, engagement of anabolic metabolism, regulation of autophagy and effector responses. Whilst Asn uptake is required for initial activation and optimal differentiation of both CD4^+^ and CD8^+^ T cells to an inflammatory cytokine-producing effector phenotype, a key finding of this work was that CD8^+^ T cell activation was less impacted by Asn deprivation as compared to CD4^+^ T cells, with this resilience being associated with elevated expression of ASNS by CD8^+^ effector cells. Activation of CD4^+^ T cells (the current work) and CD8^+^ T cells [15] in the absence of Asn is dependent upon ASNS expression, which is upregulated during activation and endows effector cells with the capacity to function independently of Asn availability. Previous work determined that TCR-induced ASNS expression is mediated via Myc-dependent pathways [15]. Myc expression is elevated in TCR-stimulated CD8^+^ T cells as compared to CD4^+^ T cell counterparts [4], consistent with higher levels of ASNS expression and enhanced capacity for activation under condition of Asn deprivation by this T cell subset. Furthermore, the biomass and protein content of TCR-stimulated CD4^+^ T cells is lower than that of CD8^+^ T cells [4; 25], further emphasizing differences in the extent of activation-induced upregulation of protein synthesis pathways between the two major T cell subsets.

Previous studies and the current work are in broad agreement regarding the importance of Asn uptake in initial T cell activation [15; 16; 17; 20; 21; 22; 23]. By contrast, recent studies have reported that Asn-deprivation can have beneficial effects on longer-term effector T cell metabolic fitness and *in vivo* anti-tumour responses [20; 21; 26]. Similarly, engineering CAR-T cells to overexpress asparaginase has been reported to enhance their function by favouring central memory phenotypes [27]. In our studies, we have shown that effector T cell responses are initially impaired in the absence of Asn but that, concomitant with elevated ASNS expression, both CD4^+^ and CD8^+^ T cells overcome these defects (present work and [15]). However, we have not replicated the finding that Asn-deprivation results in enhanced effector T cell responses, at up to 6 days after T cell stimulation [15]. Whilst our *in vitro* studies do not replicate the full complexity of *in vivo* immune responses, previous work determined that dietary Asn restriction impeded T cell responses to *Listeria* infection *in vivo* [16] whilst bacterial asparaginase expression impedes effective T cell anti-tumour responses [23; 28]. Indeed, recent evidence indicates that gut microbiota asparaginase expression may be linked to colorectal cancer progression or development [28]. Therefore, it appears that Asn restriction has context-and model-dependent impacts on T cell responses both *in vitro* and *in vivo*.

A pertinent question arising from the current and previous studies is: how do naïve T cells that express very low levels of ASNS upregulate Myc and subsequently ASNS under conditions of Asn-deprivation? It is possible that high autophagic flux, maintained after TCR-stimulation under conditions of Asn-deprivation but not in Asn-replete conditions, supplies sufficient amino acids to enable Myc expression. It appears that Asn restriction raises the threshold for T cell activation, with fewer cells reaching that threshold after stimulation. Notably, those T cells that do overcome the activation threshold in Asn-deprived conditions display similar levels of ATP production as cells activated in control conditions. Furthermore, effector CD4^+^ Th1 cells and CD8^+^ effector CTLs, generated under Asn-replete conditions, did not reintroduce autophagy upon transfer to Asn-depleted conditions. This strongly implies that ASNS expression in effector T cells negates the need for autophagy under Asn-deprivation conditions, but this will require further investigation to test directly. Notably, the regulation of T cell autophagic flux by Asn availability differs from the effects of other non-essential / conditionally-essential amino acids such as glutamine and arginine. Thus, deprivation of either glutamine and arginine prevents autophagy repression both during early T cell activation and in effector T cells [13], whereas Asn-deprivation selectively impacts autophagy in early activation.

In summary, the present works adds further to our understanding of the requirements for Asn uptake and biosynthesis in T cell activation. We provide evidence that CD4^+^ and CD8^+^ T cells display differences in their Asn requirements, and demonstrate how Asn availability impacts T cell activation, effector cytokine production, metabolic reprogramming and autophagy. Given the exciting findings demonstrating potential therapeutic roles for manipulation of T cell Asn metabolism [17; 20; 21; 23], future work will continue to investigate how manipulation of Asn availability or synthesis can impact T cell responses *in vivo*.

## Methods

### Mice

C57BL/6 and *Asns*^Tm1a^ [15] gene-trap mice were maintained at the University of Leeds St James’s Biomedical Services (SBS). Autophagy reporter mice express an mCherry-GFP-Map1lc3b (mCherry-GFP-LC3b) fusion protein under the control of the ubiquitously expressed *ROSA26* locus [29] and were maintained at the University of Dundee Biological Resource Unit. All experiments were performed according to UK Home Office regulations and the ARRIVE 2.0 guidelines, using age-and sex-matched mice (7-12 weeks). Mice were group-housed according to litter size with food and water available ad libitum, on a 12:12h light: dark schedule and maintained at constant temperature (20±1o C) and humidity.

### Cell culture

T cells were obtained from mouse lymph nodes (LN). Single cell suspensions were obtained by dissociating LN through a 70 μm filter. In some experiments, naïve CD4^+^ T cells were purified by magnetic sorting (Miltenyi Biotech). Cells were cultured in DMEM medium (Gibco) containing 10 mM glucose, supplemented with 5% dialysed FBS (Gibco), penicillin-streptomycin (Gibco), β-mercaptoethanol (50 μM, Gibco) ± asparagine (Sigma Aldrich) ± glutamine (Gibco), at concentrations stated in individual experiments at 37°C / 5% CO_2_. CD4^+^ T cells and LN cells were activated with plate-bound (purified cells) or in-solution (LN cells) CD3 (1 μg/mL, Biolegend) and CD28 mAbs (3 μg/mL, Biolegend) plus recombinant mouse IL-12 (10 ng/mL, Peprotech). For assessment of effector cytokine production, day 3 activated LN T cells were re-stimulated with phorbol 12, 13-dibutyrate and ionomycin for 3h in the presence of Brefeldin A (2.5 μg/mL, Tocris). As a positive control (amino acid-free conditions) for reintroduction of autophagic flux in effector T cells, day 5 activated cells were transferred to Hanks Balanced Salt Solution (Gibco) supplemented with dialysed FBS, penicillin-streptomycin and β-mercaptoethanol.

### Flow cytometry

The following antibodies were used: CD8β-phycoerythrin cyanine 7 (PE-Cy7) (clone YTS156.7.7, BioLegend), CD4-allophycocyanin (APC) (clone GK1.5, BioLegend), CD71-fluorescein isothiocyanate (FITC) (clone R17217, BioLegend), IFNγ-alexa fluor 488 (AF488) (clone XMG1.2, BioLegend), Tbet-PE (clone 4B10, BioLegend), TNF-peridin chlorophyll protein cyanine5.5 (PerCP-Cy5.5) (clone MP6-XT22, BioLegend), ASNS-AF647 (clone E6C2C, Cell Signaling Technology). For assessment of intracellular markers, cells were fixed and permeabilised using FoxP3 fix/perm buffer (eBioscience/ThermoFisher). For live/dead discrimination, LD Aqua (Life Technologies) was used. For analysis of mitochondrial ATP production, BioTracker^TM^-ATP-Red Live Cell Dye (5 μM, Sigma Aldrich) was added to cell cultures for 15 minutes prior to surface staining. Flow cytometry was performed using Cytoflex S (Beckman Coulter) or LSR Fortessa (BD Biosciences) flow cytometers and analysed using Flowjo 10 software (Becton Dickinson).

### ELISA

For IL-2 secretion experiments, cells were incubated with CD25 blocking Ab (Biolegend) to prevent IL-2 consumption. Supernatants were collected after 24h of culture and stored at-20°C, until analysis. Supernatant IL-2 concentrations were determined using Mouse Duoset IL-2 ELISA (R&D Systems), following manufacturer’s instructions.

### Autophagic flux experiments

Analysis of T cell autophagic flux was performed using LN T cells from mCherry-GFP-LC3b expressing reporter mice, as described previously [13]. Briefly, autophagic flux is quantified by analysis of GFP fluorescence quenching relative to mCherry signals, as assessed by flow cytometry. The mCherry/GFP ratio is measured as a “derived parameter” using Flowjo software.

### Statistics

Statistical significance (*p* value < 0.05) was determined by performing one-or two-way ANOVA with Tukey’s multiple comparisons tests, or unpaired Student’s t-tests using GraphPad Prism software, as described in figure legends. Dots in graphs represent replicate samples within an individual experiment and error bars represent SD, unless otherwise described.

### Study approval

Mouse breeding and experiments performed at the Universities of Leeds (UoL) and Dundee (UoD) were reviewed by the UoL or UoD Animal Welfare and Ethical Review Boards and all regulated work was approved by and subject to the conditions of UK Home Office Project Licences PP9249234 (UoL) and P4BD0CE74 (UoD).

## Data availability

The datasets generated during and/or analysed during the current study are available from the corresponding author on reasonable request.

## Author contributions

MS and RS designed the study. MS, LS, MT and RS acquired and analysed the data. ML and RS supervised the project. RS wrote the first draft of the manuscript. All authors, read and discussed manuscript drafts.

## Supporting information

All supplemental figures

