## Supplementary material for "Asparagine availability differentially regulates early vs late CD4^+^ and CD8^+^ T cell activation, metabolism and autophagy": All supplemental figures

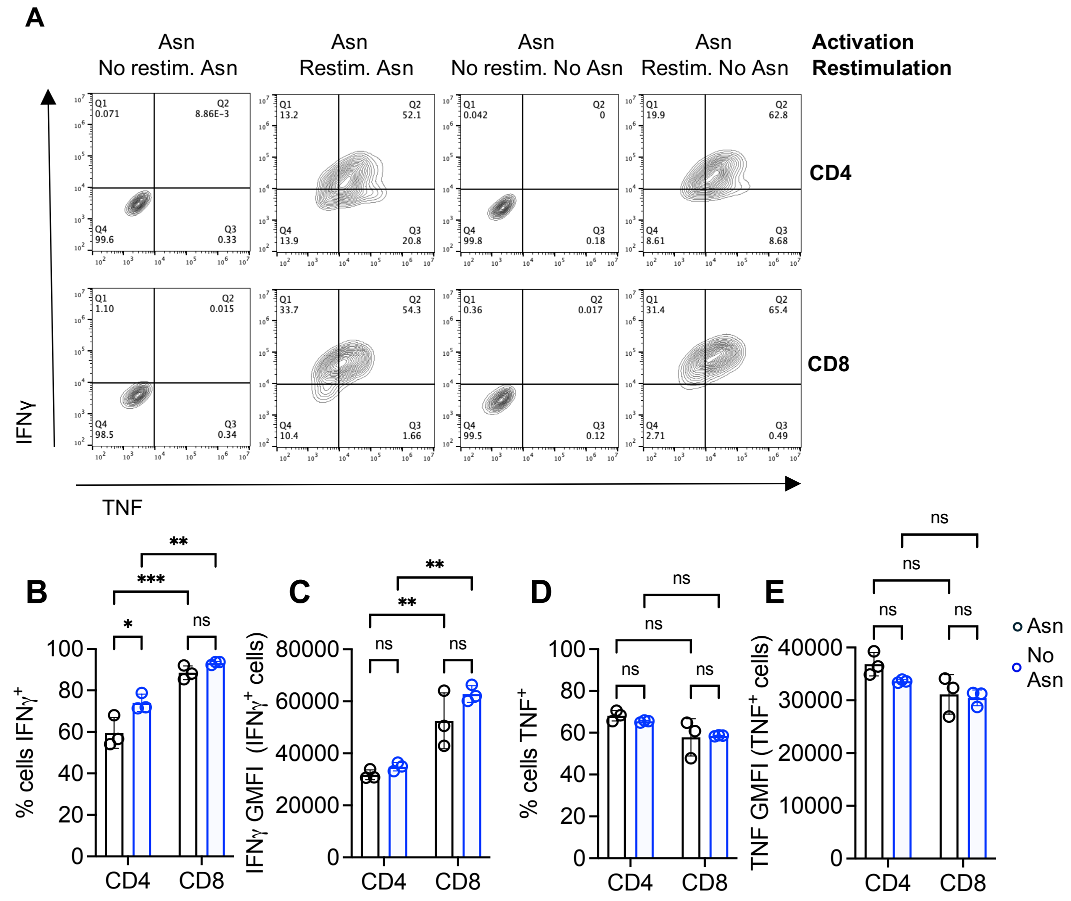


**Supplementary Figure 1.** Asn availability is not required for cytokine production in effector T cells. To assess requirements for Asn in effector T cell cytokine production, lymph node T cells were differentiated to a Type 1 effector phenotype by stimulating for 72h with CD3 / CD28 antibodies and IL-12 in Asn-replete media, then restimulated with PDBU/Ionomycin in DMEM supplemented ± Asn. Representative dotplots (A) show levels of IFNγ and TNF staining in gated $CD4^+$ and $CD8^+$ T cells, whilst barcharts show mean proportions of $IFN\gamma^+$ (B) and $TNF^+$ (D) cells or geometric mean fluorescence intensity (GMFI) or cytokine staining in gated cytokines-positive cells (C, E). Data are from 1 of 3 repeated experiments. In all cases, error bars represent SD. ns – not significant, * p<0.05, ** p<0.01, *** p<0.001, **** p<0.0001 as determined by two-way ANOVA with Tukey's multiple comparisons test.

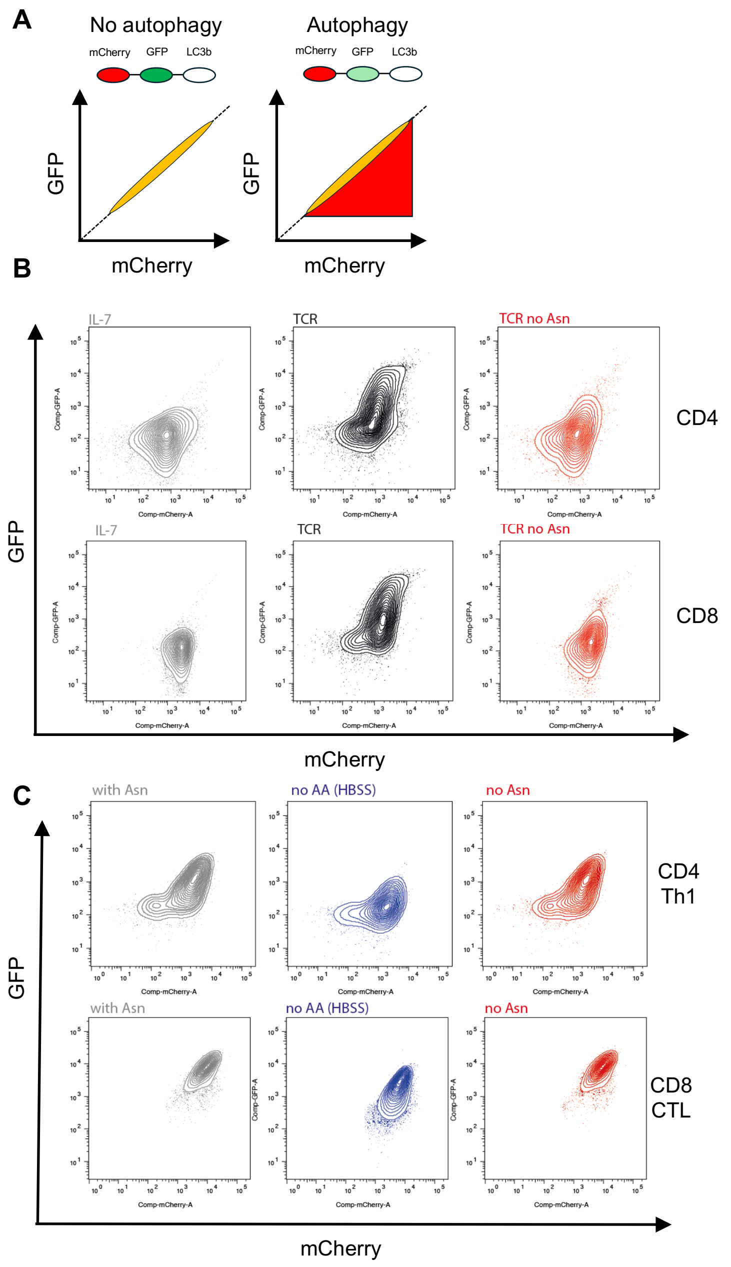


**Supplementary Figure 2.** Differential roles for Asn availability in regulation of autophagic flux in naïve vs effector T cells. (A) Levels of autophagic flux were assessed using mCherry-GFP-LC3b reporter mice by analysis of relative levels of GFP and mCherry signal: linear GFP-mCherry represent autophagy repression, quenching of GFP signal reflects recruitment of LC3b to autolysosomes and high autophagic flux. (B) Representative dotplots show levels of GFP and mCherry fluorescence in gated $CD4^+$ and $CD8^+$ T cells activated for 18h ± 300 μM Asn. (C) Effector T cells from autophagy reporter mice were generated by 5 days of stimulation *in vitro*. Cells were switched into DMEM ± Asn or amino acid-free conditions (HBSS) for 2h followed by assessment of autophagy.

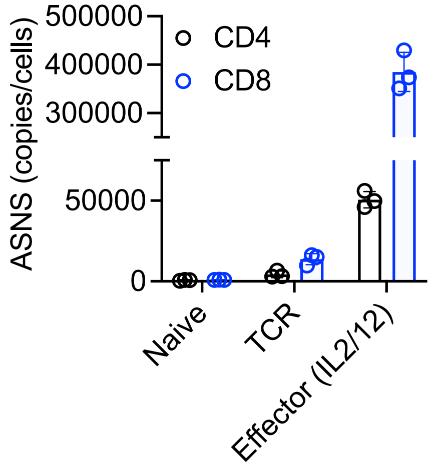


**Supplementary Figure 3.** Increased expression of ASNS in effector CD8$^+$ as compared to CD4$^+$ T cells. Data are taken from the Immunological Proteome Resource (Ref. 25) and represent biological replicate values of ASNS protein copies / cell.
